# Mechanical confinement governs phenotypic plasticity in melanoma

**DOI:** 10.1101/2024.01.30.577120

**Authors:** Miranda V. Hunter, Emily Montal, Yilun Ma, Reuben Moncada, Itai Yanai, Richard P. Koche, Richard M. White

## Abstract

Phenotype switching is a form of cellular plasticity in which cancer cells reversibly move between two opposite extremes - proliferative versus invasive states. While it has long been hypothesised that such switching is triggered by external cues, the identity of these cues has remained elusive. Here, we demonstrate that mechanical confinement mediates phenotype switching through chromatin remodelling. Using a zebrafish model of melanoma coupled with human samples, we profiled tumor cells at the interface between the tumor and surrounding microenvironment. Morphological analysis of these rare cells showed flattened, elliptical nuclei suggestive of mechanical confinement by adjacent tissue. Spatial and single-cell transcriptomics demonstrated that the interface cells adopted a gene program of neuronal invasion, including acquisition of an acetylated tubulin cage that protects the nucleus during migration. We identified the DNA-bending protein HMGB2 as a confinement-induced mediator of the neuronal state. HMGB2 is upregulated in confined cells, and quantitative modelling revealed that confinement prolongs contact time between HMGB2 and chromatin, leading to changes in chromatin configuration that favor the neuronal phenotype. Genetic disruption of HMGB2 showed that it regulates the trade-off between proliferative and invasive states, in which confined HMGB2^high^ tumor cells are less proliferative but more drug resistant. Our results implicate the mechanical microenvironment as a mechanism driving phenotype switching in melanoma.

## INTRODUCTION

The ability of cancer cells to adopt new phenotypes without additional DNA mutations is now well understood to significantly influence tumor behavior and aggressiveness^1^. Such plasticity has long been observed in melanoma, where early studies identified transcriptomic and phenotypic states not linked to a particular genetic lesion^2–4^. The advent of single-cell RNA-sequencing has provided further nuance to these observations, suggesting that most tumors are composed of a heterogeneous, yet reproducible number of cell states, referred to as meta-programs^5–7^. The extent to which tumor cells transition between states is an open area of investigation that has been hypothesized to be triggered by unknown cues from the local microenvironment. The identification of such cues has clinical relevance, as they may enable the conversion of a superficial melanoma into an invasive and drug resistant one^8–10^. Here, using a combination of zebrafish transgenics and human samples, we show that mechanical confinement from the surrounding microenvironment induces stable changes in chromatin architecture that cause the tumor cell to transition from a proliferative to invasive state.

## RESULTS

### Tumor gene expression is influenced by the local microenvironment

To study the influence of the local microenvironment on tumor invasion, we applied spatially resolved transcriptomics and single-cell RNA-sequencing (scRNA-seq) to a transgenic zebrafish model of *BRAF^V600E^*-driven melanoma (**Fig. 1a**). Tumors from this model frequently invade into adjacent tissues, including the underlying dermis and muscle. Across fish, we found a conserved “interface” transcriptional cell state that occurs where tumor cells invade into the microenvironment^11^ (**Fig. 1a**).

**Figure 1:**
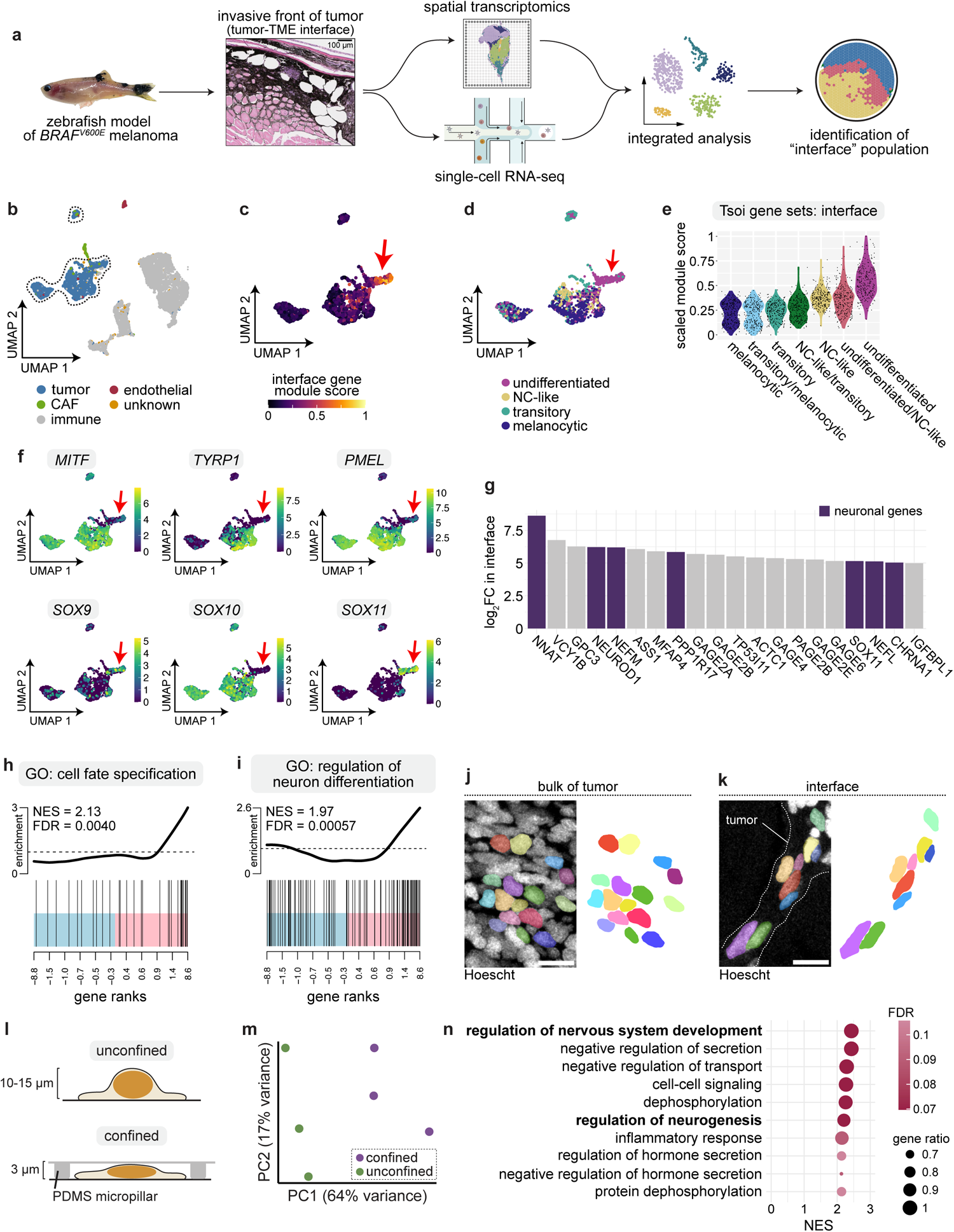
Confinement induces an undifferentiated neuronal gene program. **a.** Schematic detailing workflow of spatial transcriptomics and single-cell RNA-seq experiments performed on zebrafish melanomas. **b.** UMAP of human melanoma scRNA-seq dataset from Jerby-Arnon et al. Cluster annotations from the original manuscript are labeled. Tumor cell clusters are outlined. **c.** Gene module scoring for interface genes extracted from zebrafish spatial transcriptomics and scRNA-seq data, projected onto tumor cells outlined in b. Red arrow denotes the subpopulation with highest expression of interface genes. **d.** Cell state classification for melanoma differentiation states identified by Tsoi et al. Cells were classified based on highest expression of the gene modules indicated. **e.** Module scores for melanoma cell state genes from Tsoi et al. in interface cells. **f.** Normalized expression per cell in UMAP space for the indicated genes. Red arrow indicates the interface cluster identified in b. **g.** Top 20 most highly upregulated genes in the human interface cluster. Neuronal genes are labeled in purple. **h-i.** GSEA barcode plot for the indicated pathways. Normalized enrichment score (NES) and false discovery rate (FDR) are labeled. **j-k.** Immunofluorescence of adult zebrafish tissue sections highlighting the center of the tumor (j) and tumor-TME interface (k). Individual nuclei are pseudocoloured and displayed without image overlay at right. Scale bar, 10 µm. **l.** Schematic of in vitro confinement workflow. **m.** Principal component analysis plot for each RNA-seq replicate. Percent variance for each principal component is labeled. **n.** Top 10 most highly upregulated pathways from GSEA of confined cells relative to unconfined. NES and FDR are indicated. Neuronal pathways are highlighted.

To investigate if these interface cells occur in human patients, we analyzed a recently published human melanoma scRNA-seq dataset, composed of 7,186 tumor and stromal cells from 31 patients with either untreated or immunotherapy-resistant melanoma^12^ (**Fig. 1b**). We scored every tumor cell in the dataset for the relative expression of interface marker genes extracted from our zebrafish transcriptomics datasets. Similar to our previous observations^11^, a subpopulation of human tumor cells highly upregulated interface genes identified from our zebrafish dataset (**Fig. 1c**). To better understand the nature of these cells, we compared our interface population to human melanoma cell states. Recent work defined at least 4 cell states that encompass the differentiation trajectory of the melanoma cell (melanocytic, transitory, neural crest-like, undifferentiated)^13^. We scored tumor and interface cells for the relative expression of gene modules encompassing the entire melanoma cell differentiation trajectory, and classified cells into each of the 4 states based on their expression of gene modules annotated for that state^13^. While the bulk of the tumor cells appeared to be relatively evenly distributed between the 4 cell states (**Fig. 1d**), interface cells displayed a clear upregulation of genes characteristic of the undifferentiated state (**Fig. 1d-e**).

Previous work has shown that melanoma cell behavior is regulated by phenotype switching, transcriptional programs that mediate transitions between differentiated/proliferative and undifferentiated/invasive states^8,9,13,14^. The transition between the differentiated/proliferative and undifferentiated/invasive states is regulated by the transcription factors *MITF*, *SOX9*, and *SOX10*, among others^14^. *MITF* and *SOX10* have key roles in melanocyte differentiation and proliferation^15,16^, whereas SOX9 is associated with a switch from a melanocytic to an undifferentiated invasive state^17–19^. We found that *MITF* and *SOX10* were highly downregulated in human interface cells, whereas *SOX9* was highly upregulated (**Fig. 1f**). Classical melanocyte pigmentation genes, indicative of a differentiated state, were also minimally expressed in interface cells relative to the rest of the tumor, including *MITF* (log_2_FC = −5.19), *TYRP1* (log_2_FC = −7.75) and *PMEL* (log_2_FC = −8.78) (**Fig. 1f** and **Table S1**). In addition to loss of melanocyte differentiation markers in the interface cells, we also unexpectedly found an enrichment of genes involved in neuronal development, including *SOX11* (log2FC = 5.15)*, NNAT* (neuronatin, log_2_FC = 8.61), *NEUROD1* (neuronal differentiation 1, log_2_FC = 6.22), and *NEFM* (neurofilament medium chain, log_2_FC = 6.19) (**Fig. 1g** and **Table S1**). Gene set enrichment analysis (GSEA) revealed that transcriptional programs linked to cell fate specification and neuronal development were highly enriched in interface cells (**Fig. 1h-i**). These data indicate that the interface cells adopted an invasive state with markers of neuronal development.

### Confinement induces melanoma dedifferentiation towards a neuronal state

To examine factors within the local TME that may drive the interface state, we performed histology on tissue sections from our transgenic zebrafish melanoma model, focusing on the invasive front. Tumor cells invading into the TME displayed flattened, elongated nuclei when compared to cells in the bulk tumor mass (**Fig. 1j-k**). Recent work showed that tumor nuclei become highly elongated in order to squeeze through mechanically-restrictive environments^20^. While it is likely that numerous factors (secreted factors, immune infiltration, extracellular matrix stiffness) influence tumor invasion, we hypothesized that the mechanical forces exerted on the cell and nucleus during invasion may cause stable changes in gene expression and tumor cell behavior.

To test this hypothesis, we adopted a system to confine human melanoma cells (A375 cell line) *in vitro* at predefined heights (3 µm), using a PDMS piston and micropatterned coverslips^21^ (**Fig. 1l**). We confirmed that *in vitro* confinement of A375 cells does not cause cell death, as confined cells did not upregulate the apoptosis markers cleaved caspase-3, annexin V, or cleaved PARP (**Fig. S1a-c**). After removing confinement, cells recovered their typical morphology within 24 hours with no evidence of widespread cell death (**Fig. S1d**). To define the transcriptional changes induced by mechanical force, we confined A375 cells for ∼18 hours and performed bulk RNA-sequencing, comparing gene expression to unconfined cells (**Fig. 1l**). Principal component analysis revealed a significant amount of transcriptional alterations induced by confinement (**Fig. 1m** and **Table S2**). Similar to what we observed in interface cells from human patients, many neuronal pathways were upregulated in confined cells, including regulation of nervous system development and neurogenesis (**Fig. 1n** and **Table S2**). Together, these data suggest that mechanical confinement causes melanoma cells to adopt a neuronal identity.

### The microtubule cytoskeleton hijacks neuronal mechanisms to reinforce the tumor cell against mechanical force

Neuronal development is heavily reliant on microtubule (MT) architecture, with the MT cytoskeleton influencing almost every aspect of neuronal structure and function^22^. Importantly, the cytoskeleton is the main mechanism by which mechanical force is transmitted within and between cells, and often remodels in response to mechanical stimuli^23^. Recent reports indicated the MT cytoskeleton is stabilized by force to protect confined cells from damage^24–27^. We hypothesized that melanoma cells hijack neuronal mechanisms to allow them to invade into the mechanically confined microenvironment. Using our system to apply mechanical force *in vitro*, we characterized how the MT cytoskeleton responds to confinement in tumor cells. We performed live imaging of A375 cells in which the endogenous microtubule cytoskeleton was labeled with the fluorescent dye SiR-tubulin^28^. There was significant reorganization of the MT cytoskeleton in confined cells: while the MT cytoskeleton initially resembled that of unconfined cells, with a central microtubule organizing center (MTOC) from which linear MTs radiated outward, in most confined cells within 2-4 hours curved MTs instead began encircling both the cell and nuclear periphery (**Fig. 2a-b**), reminiscent of recent work showing curved MTs represent MT stabilization against compressive forces, where MTs bend to prevent buckling or rupture^24,29^. The loss of a central MTOC and radial microtubules we observed in confined cells is reminiscent of that of neurons, in which microtubule organization is typically centrosome-independent^22,30–32^. These results indicate that confined melanoma cells rapidly undergo structural changes in the MT cytoskeleton similar to that seen in neurons.

**Figure 2:**
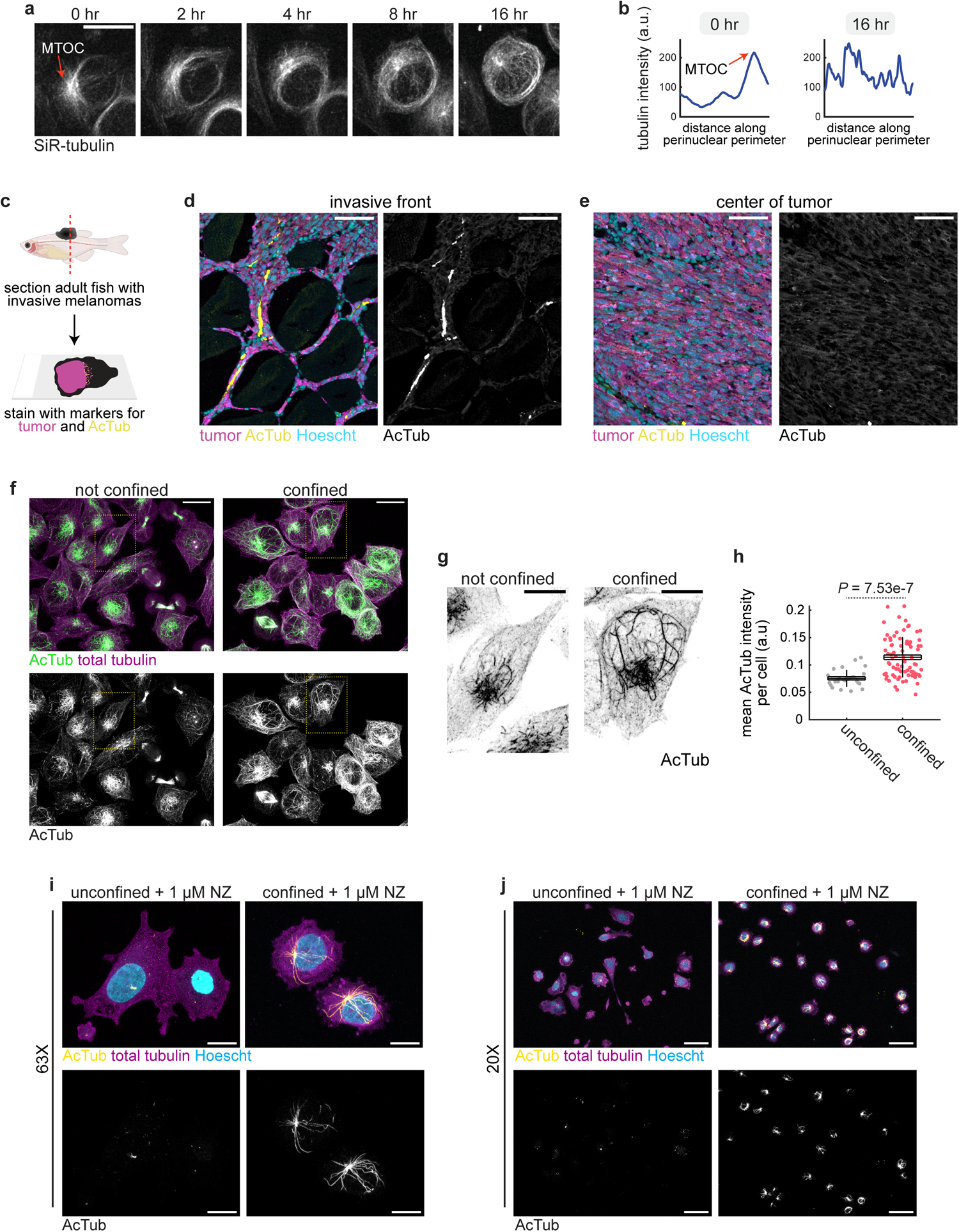
A perinuclear acetylated tubulin cage assembles in response to confinement. **a.** Representative stills from confocal imaging of A375 cells stained with SiR-tubulin. **b.** Line intensity profile of perinuclear tubulin intensity over time from the images shown in a. **a-b.** Microtubule organizing center (MTOC) is highlighted. **c.** Schematic detailing immunofluorescence staining of sections from adult zebrafish melanomas. **d-e**. Immunofluorescence images of acetylated tubulin staining at the invasive front (d) compared to the center of the tumor (e). Scale bars, 50 µm. **f.** A375 cells stained with antibodies labeling acetylated tubulin (green) and total tubulin (purple). Scale bars, 20 µm. **g.** Inset of regions labeled in f. Scale bars, 10 µm. **h.** Quantification of whole-cell acetylated tubulin intensity. Each point represents one cell. Unconfined: *n* = 27 cells from 3 images. Confined: *n* = 80 cells from 9 images. **i-j.** A375 cells treated with 1 µm nocodazole (NZ) for ∼18 hours and stained for acetylated tubulin (yellow at top, and bottom), total tubulin (purple) and Hoescht (blue). Magnification is indicated. Scale bars, 50 µm (j) and 10 µm (i).

Acetylated tubulin is a critical mediator of neuronal architecture and function, and is a widely used marker of axonal microtubules^33^. Tubulin acetylation stabilizes microtubules, and thus the cell, against mechanical pressure^27,29^. Curved microtubules such as those we observed in confined cells (**Fig. 2a-b**) are also indicative of long-lived, stabilized microtubules^29^. When we performed IF on tumor sections from adult fish with transgenic *BRAF^V600E^*-driven melanomas, we observed a significant accumulation of acetylated tubulin at the tumor border^11^ (**Fig. 2c-e**). Similarly, when we confined A375 cells *in vitro*, we quantified a significant upregulation of acetylated tubulin upon confinement (*P* = 6.86e-12; **Fig. 2f-h**). Acetylated tubulin filaments in unconfined cells were typically short and linear, whereas in confined cells acetylated tubulin filaments were longer and more curved, again indicative of stabilization (**Fig. 2f-g**).

We noticed that in confined cells, the hyperacetylated tubulin network was often perinuclear (**Fig. 2f-g**), and hypothesized this network may be providing structural support specifically to the nucleus. As the stiffest and largest organelle in the cell, the nucleus is particularly vulnerable to confinement-induced mechanical stress, with confined migration often causing nuclear envelope rupture and DNA damage^34–37^. During migration through confined spaces, neurons assemble a perinuclear network of acetylated tubulin to protect the nucleus from mechanical damage^38–41^. We confirmed the stability of the perinuclear acetylated tubulin network by treating confined A375 cells with the microtubule depolymerizing agent nocodazole (NZ). Acetylated microtubules are resistant to NZ^29^. 1 µM NZ induced disassembly of almost all non-modified tubulin filaments in both unconfined and unconfined cells (**Fig. 2i-j**). While there was little acetylated tubulin present in unconfined, NZ-treated cells, almost every confined NZ-treated cell contained a highly stable perinuclear acetylated tubulin cage enveloping the nucleus (**Fig. 2i-j**).

In addition to acetylation, microtubules exhibit a variety of post-translational modifications (PTMs), including tyrosination, glutamylation, glycylation, and methylation, many of which have been reported to influence microtubule stability^42^. To better understand the structure of the perinuclear acetylated tubulin network, we used IF to characterize the scope of microtubule PTMs in confined melanoma cells. Detyrosination has been linked to longer-lived stabilized MTs^43,44^. However, most microtubules in both confined and unconfined A375 cells were highly tyrosinated, with very few detyrosinated MTs found in both unconfined and confined cells (**Fig. S2a-d**). This is likely related to the high levels of tyrosine and tyrosinase typically found in melanoma^45^. While confined perinuclear MTs were occasionally tyrosinated and, rarely, detyrosinated (**Fig. S2a-d**), there was no specific enrichment of tyrosinated or detyrosinated MTs in the perinuclear network as we observed for acetylated MTs. Similar results were observed for polyglutamylated MTs (**Fig. S2e-f**). There was no evidence of polyglycylated MTs in both unconfined and confined cells (**Fig. S2g-h**). This suggests that acetylation is the primary PTM mediating the assembly and/or stability of the perinuclear tubulin network. Together, these data indicate that confined melanoma cells assemble a highly stable perinuclear tubulin network to reinforce the nucleus against mechanical stress, similar to that of neurons.

### HMGB2 is a confinement-induced marker of invasion

Our results so far suggest that confinement induces a neuronal identity in melanoma cells at the interface between tumor and TME, characterized by changes in cytoskeletal architecture and gene expression. We examined our zebrafish and human transcriptomics datasets to identify potential confinement-induced mediators of this state. Interface cells at the invasive front were characterized by consistent upregulation of high mobility group (HMG)-family proteins (**Fig. 3a**), which control chromatin architecture by binding to and bending DNA without sequence specificity to relieve mechanical strain^46^. While a HMG-enriched transcriptional program is consistently upregulated across a wide range of tumor types^6^, including melanoma^47^, the contribution of this program to tumor progression is not well understood. We focused on HMGB2 as it was the most highly upregulated HMG-family member in interface cells in our zebrafish transcriptomics data (zebrafish orthologs *hmgb2a* and *hmgb2b*; **Fig. 3a**). HMGB2 was also highly upregulated in interface cells in human patient data^12^ (*P* = 3.24e-37; **Fig. 3b-c**).

**Figure 3:**
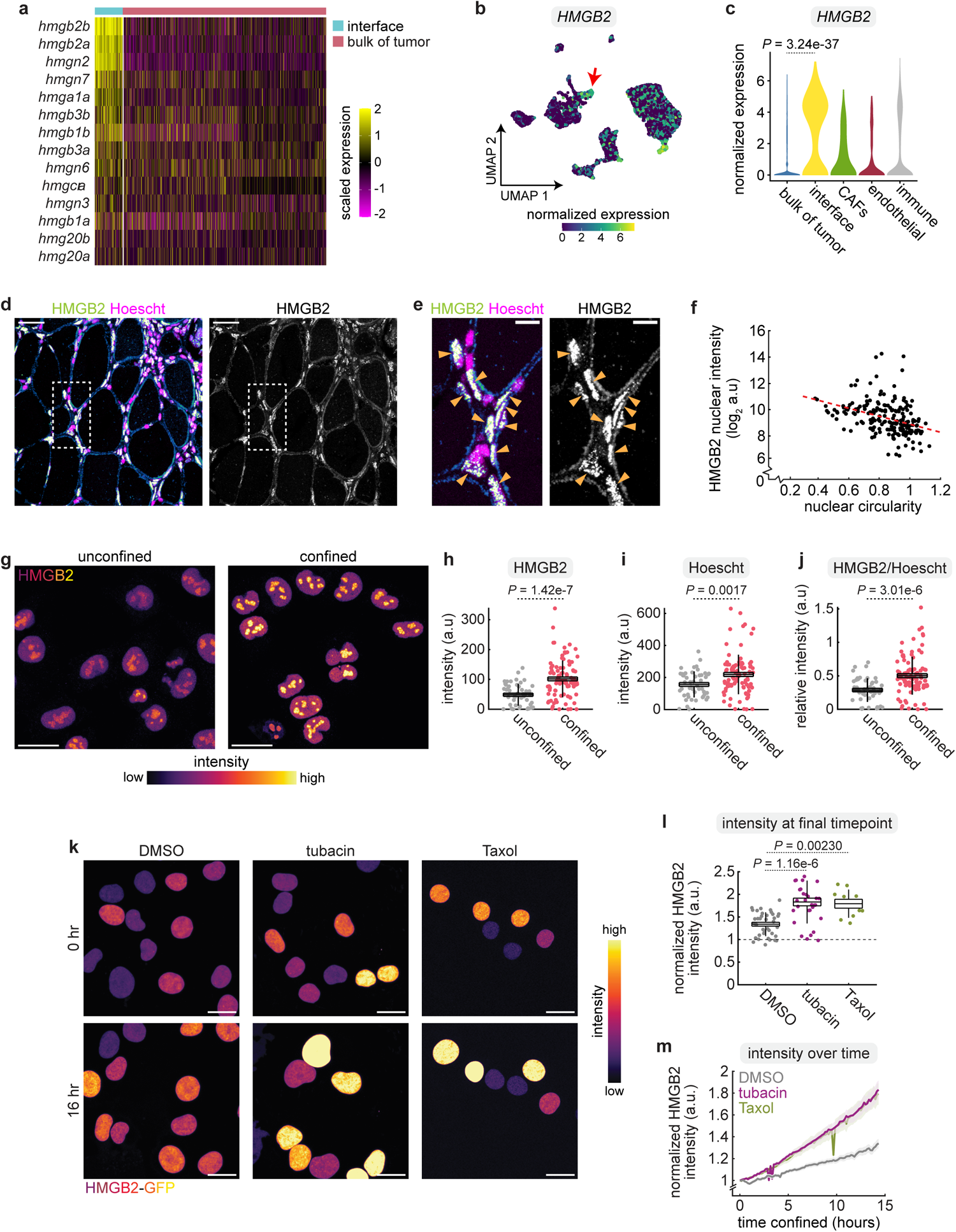
HMGB2 is a confinement-induced marker of invasion. **a.** Heatmap of HMG-family member expression in invasive tumor cells relative to the bulk of the tumor from zebrafish melanoma scRNA-seq data. **b.** *HMGB2* expression per cell in UMAP space from human melanoma scRNA-seq data from Jerby et al. Arrow indicates the interface cluster. **c.** Violin plot of mean *HMGB2* expression per cluster. *P*-value calculated using Wilcoxon rank sum test. **d.** Immunostained sections from a zebrafish melanoma tissue section stained for HMGB2 and Hoescht. Scale bar, 50 µm. **e.** Inset of region indicated in d. Scale bar, 10 µm. Elongated cells with high HMGB2 expression are labeled. **f.** Correlation between nuclear circularity and HMGB2 intensity. Red dashed line indicates line of best fit by linear regression. **g.** Immunofluorescence images of A375 cells stained with antibodies targeting HMGB2. Scale bars, 25 µm. **h-j.** Quantification of Hoescht and HMGB2 intensity per cell. Unconfined: *n* = 49 cells from 3 images. Confined: *n* = 97 cells from 9 images. **k.** HMGB2-GFP accumulation in A375 cells treated with DMSO, tubacin or Taxol. Scale bars, 25 µm. Time after applying confinement is indicated. **l.** HMGB2-GFP intensity in confined cells over time. **m.** HMGB2-GFP intensity per cell at the final time point imaged (∼16 hours). **l-m.** DMSO: *n* = 38 cells from 7 movies. Tubacin: *n* = 31 cells from 7 movies. Taxol: *n* = 10 cells from 3 movies.

To validate our transcriptomics results indicating HMGB2 is upregulated at the invasive front, we used immunostaining to examine the expression and localization of HMGB2 in tissue sections from zebrafish melanomas. In visibly invading tumor cells, HMGB2 was only upregulated in elongated or misshapen tumor cells that appeared to be under mechanical pressure due to confinement by adjacent tissues (**Fig. 3d-f**), suggesting confinement induces upregulation of HMGB2. HMGB2 levels were inversely correlated to circularity of the nucleus (*R* = −0.474; *P* = 1.19e-12; **Fig. 3f**). Similar to our *in vivo* zebrafish results, *in vitro* confinement of A375 human melanoma cells for ∼16 hours caused nuclear HMGB2 intensity to approximately double relative to the unconfined control (*P* = 1.42e-7; **Fig. 3g-h**). This increase in HMGB2 in confined cells was not solely due to changes in nuclear density: while Hoescht intensity also slightly increased upon confinement (likely due to compacting of the nucleus; *P* = 0.0017; **Fig. 3i**), when we normalized HMGB2 intensity to Hoescht intensity on a per-nucleus level, we still quantified a significant increase (*P* = 3.01e-6; **Fig. 3j**). We used time-lapse confocal microscopy to visualize and quantify the rate of HMGB2 upregulation in confined cells over time, using a stable A375 cell line expressing HMGB2-GFP. Nuclear HMGB2-GFP levels increased linearly over ∼16 hours of imaging (increasing by 0.269±0.020-fold per hour) to a final level of 1.76±0.072-fold relative to the first time point (**Fig. S3a-b**). Using IF, we also examined levels of HMG-family members HMGB1 and HMGA1, which were transcriptionally upregulated in interface cells, albeit to a lesser extent than HMGB2, and quantified no change in their expression upon confinement (**Fig. S3c-f**), suggesting that confinement-induced upregulation of HMGB2 is not a general property of all HMG-family members.

### HMGB2 upregulation is mediated by the perinuclear acetylated tubulin network

To investigate how mechanical confinement induces HMGB2 upregulation, we focused on the MT cytoskeleton due to its role in mechanotransduction^24–27^, hypothesizing that the perinuclear acetylated tubulin network (**Fig. 2f-g**) may function to propagate confinement-induced mechanical force to the nucleus. We thus used pharmacological inhibitors to modulate tubulin dynamics and acetylation state in confined cells to study whether the acetylated tubulin network influences HMGB2 upregulation. Tubulin acetylation is mediated by the highly conserved acetyltransferase ATAT1^48^, with deacetylation controlled by HDAC6^49^. To induce tubulin hyperacetylation, we treated A375 cells with the highly specific HDAC6 inhibitor tubacin^50^, and confirmed that tubacin increased tubulin acetylation without affecting histone acetylation (**Fig. S4**). In confined cells, tubacin treatment significantly increased the total fold change of nuclear HMGB2 accumulation (*P* = 1.1646e-6; **Fig. 3k-l**), as well as increasing the rate of accumulation, which approximately doubled relative to the vehicle control (0.353±0.0364-fold per hour vs 0.145±0.0171-fold per hour; *P* = 1.1589e-6; **Fig. 3m**). To stabilize MTs in an acetylation-independent manner, we treated confined cells with 100 nM paclitaxel (Taxol). Taxol stabilizes microtubules by binding β-tubulin and preventing incorporation of tubulin monomers into the MT filament^51^. Taxol treatment significantly increased HMGB2 accumulation in confined cells (*P* = 0.00230, **Fig. 3k-l**). The rate of HMGB2 accumulation upon confinement was remarkably similar between Taxol- and tubacin-treated cells (0.338±0.0425-fold per hour vs 0.353±0.0364-fold per hour; *P* = 0.961; **Fig. 3m**), indicating HMGB2 enrichment upon confinement is linked to microtubule stability, rather than the acetylation state itself or other functions for HDAC6.

To confirm a role for the perinuclear acetylated tubulin network in mediating HMGB2 accumulation in confined cells, we treated A375[HMGB2-GFP] cells with 1 µM nocodazole (NZ) immediately before applying confinement. Since the perinuclear acetylated tubulin cage is resistant to NZ (**Fig. 2i-j**), NZ treatment allows us to specifically interrogate the contribution of acetylated MTs to HMGB2 upregulation in confined cells. While we observed an almost total loss of the visible SiR-tubulin signal in confined, NZ-treated cells as expected (**Fig. S5a**), HMGB2-GFP accumulation was unaffected (*P* = 0.714, **Fig. S5a-c**). Together, our results suggest a highly stable perinuclear acetylated tubulin network assembles in response to confinement and signals to the nucleus to trigger upregulation of HMGB2.

### Confinement stabilizes interactions of HMGB2 with chromatin

Our results so far indicate a perinuclear network of acetylated tubulin may stabilize the nucleus against mechanical stress and trigger upregulation of the chromatin modifier HMGB2. HMGB2 typically binds chromatin without sequence specificity and induces large (45°) bends in DNA to provide space for transcription factor binding and relieve mechanical strain^52,53^. To investigate whether confinement changes how HMGB2 interacts with chromatin, we used fluorescence recovery after photobleaching^54^ (FRAP) to characterize the dynamics of nuclear HMGB2 upon confinement. A375[HMGB2-GFP] cells were confined for ∼18 hours before performing FRAP on cells while still in the confinement chamber. Similar to previous reports^55,56^, HMGB2-GFP was highly dynamic even in unconfined cells (**Fig. 4a-d**), where only ∼25% of protein was stably bound within the nucleus (mobile fraction = 75.21±0.91%), supporting a model in which chromatin-associated proteins exhibit rapid 3D diffusion and stochastic interactions with DNA in order to find their specific binding site^57,58^. In both unconfined and confined cells, the FRAP recovery curves fit well to a 2-component exponential equation (average *R^2^* = 0.986±0.00054; **Fig. 4d-e**; see Methods), suggesting presence of 2 pools of HMGB2 within the nucleus: a fast-diffusing pool representing random interactions with chromatin, and a slower-diffusing pool representing more specific, stable interactions^59^. We used the fitted 2-component exponential to calculate the relative proportion of fast- and slower-diffusing HMGB2. In both confined and unconfined cells, most (>90%) of HMGB2 protein was fast-diffusing. However, confinement significantly increased the proportion of slower-diffusing HMGB2 (6.76±0.39% vs. 4.21±0.39%; *P* = 1.134e-5; **Fig. 4f**), suggesting that in confined cells, HMGB2 exhibits more specific, stable interactions with chromatin.

**Figure 4:**
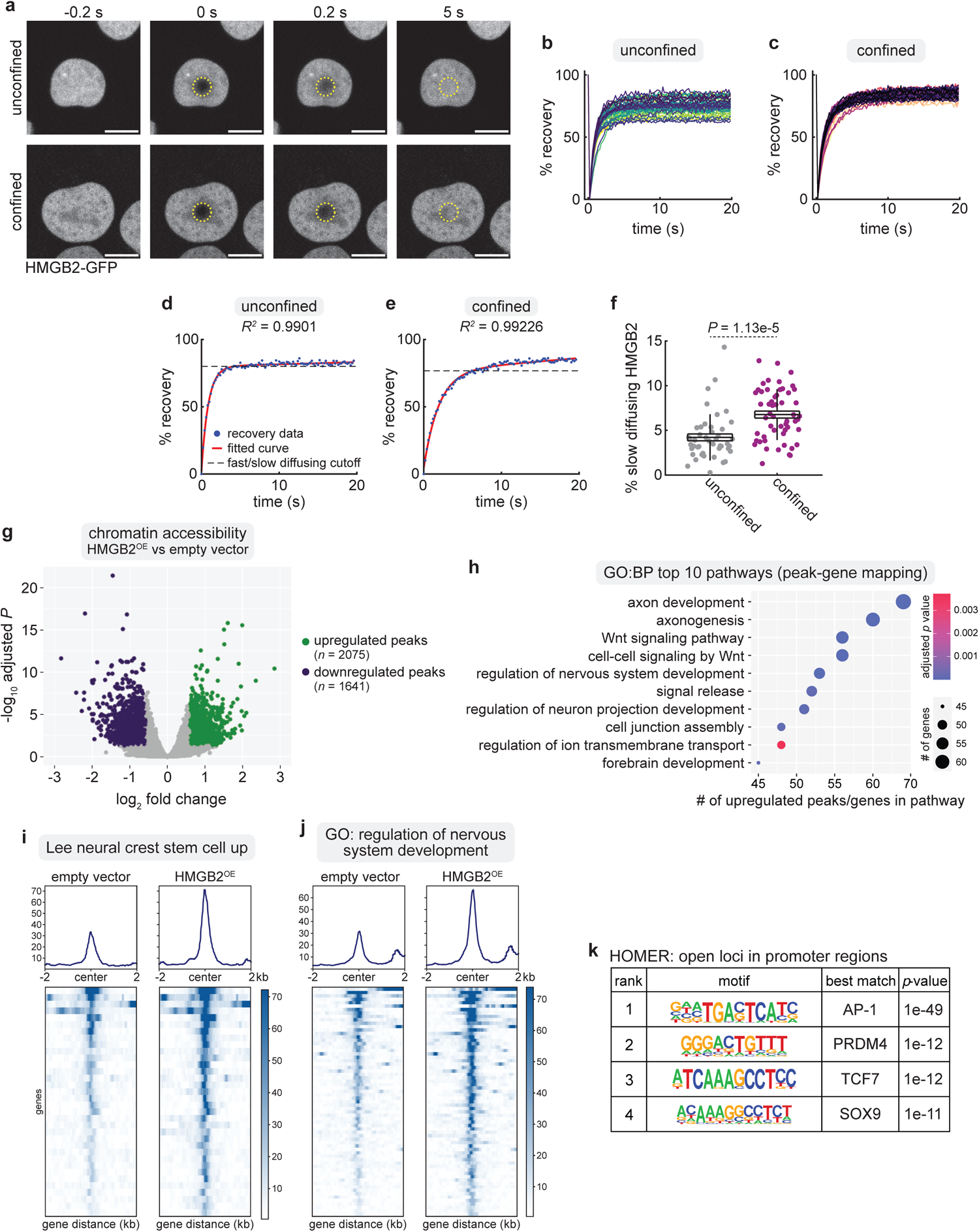
Confinement-mediated stabilization of HMGB2 increases chromatin accessibility at neuronal loci. **a.** Representative stills from timelapse imaging of A375 cells expressing HMGB2-GFP and subjected to FRAP. Yellow dashed region indicates photobleached area. Time is relative to photobleaching. **b-c.** FRAP recovery curves for HMGB2-GFP in unconfined (b) and confined (c) cells. Each curve represents fluorescence recovery within area photobleached on a single cell. **d-e.** Representative plots showing a 2-component exponential equation fit to HMGB2-GFP fluorescence recovery curves in unconfined (d) and confined (e) cells. **f.** Relative proportion of slow-diffusing HMGB2-GFP. **b-e**, **f.** Unconfined: *n* = 45 cells. Confined: *n* = 54 cells. **g.** Volcano plot of differentially expressed peaks upon HMGB2 overexpression. *P*-value cutoff, 0.05; fold change cutoff, log_2_(1.25) and log_2_(−1.25). **h.** Top 10 enriched pathways from genes mapped to open chromatin loci upon HMGB2^OE^. **i-j.** Tornado plots showing chromatin accessibility at loci linked to genes in the indicated pathways. **k.** HOMER de novo motif analysis of transcription factor motifs in open chromatin regions in promoter regions of genes upon HMGB2^OE^.

### HMGB2 increases chromatin accessibility at neuronal loci

As our FRAP analysis suggests mechanical force upregulates HMGB2 and stabilizes its interactions with chromatin, we hypothesized this would likely influence chromatin accessibility. To test this, we performed ATAC-seq on A375 cells overexpressing HMGB2. Overexpression of HMGB2 broadly increased chromatin accessibility (**Fig. 4g**). When we mapped open peaks to genes and performed GO pathway analysis, we found that regions of open chromatin were highly enriched for neuronal genes (**Fig. 4h** and **Table S3**). Upregulation of HMGB2 increased chromatin accessibility at loci linked to neural crest and neuronal development (**Fig. 4i-j** and **Table S3**). We used HOMER *de novo* motif analysis^60^ to identify conserved transcription factor binding motifs enriched within the promoter of open chromatin regions upon HMGB2^OE^. The top-ranked motif was an *AP-1* motif (**Fig. 4k**; *P* = 1e-49), which is linked to melanocyte reprogramming^61^ and melanoma plasticity^62^. Other highly ranked motifs included *PRDM4*, which functions in neural development^63–65^ (**Fig. 4k**; *P* = 1e-12), and *SOX9*, which is a critical regulator of the pro-invasive phenotype switch in melanoma^17^ (**Fig. 4k**; *P* = 1e-11). Together, these data indicate that upregulation of HMGB2 increases chromatin accessibility specifically at neuronal loci, promoting an overall confinement-induced dedifferentiation program.

To examine this at the transcriptional level, we performed bulk RNA-seq on A375 cells stably overexpressing HMGB2 (**Fig. S6a**). HMGB2^OE^ A375 cells appeared to adopt a mesenchymal-like, invasive morphology relative to the empty vector control (**Fig. S6b**). The most highly upregulated gene upon HMGB2 overexpression was *UNC5D*, a netrin receptor that promotes neuronal survival and migration^66–68^ (**Fig. S6c** and **Table S4**). A number of other neuronal genes were also highly upregulated, including *DCC*, *SH3GL2*, and *NPX2* (**Fig. S6c-d** and **Table S4**). GSEA revealed an overall overrepresentation of neuronal gene programs within enriched pathways (**Fig. S6e-f** and **Table S4**). In support of a pro-invasive role for HMGB2 in melanoma, a number of invasive genes were also upregulated upon HMGB2 overexpression (*MAGEA1*, *SPP1*, *CTAG2, GDF6*; **Fig. S6d** and **Table S4**). Together, these data indicate that confinement prolongs contact time between HMGB2 and chromatin to induce an invasive neuronal state.

### Confinement-induced HMGB2 mediates a tradeoff between proliferation and invasion

One prediction of the phenotype switching model in melanoma is that tumor cells can reversibly move between opposite extremes, in which there is a tradeoff: cells which are highly proliferative are less invasive, whereas cells that are highly invasive are less proliferative (**Fig. 5a**). Our data so far suggest that confinement-induced upregulation of HMGB2 could be one factor mediating the switch between these extremes. Consistent with this idea, we found that many proliferation-related pathways were downregulated in our RNA-seq data from HMGB2^high^ confined A375 cells (**Fig. 5b**). To directly visualize the influence of confinement on this tradeoff, we generated an A375 cell line stably expressing the FastFUCCI cell cycle sensor^69,70^. The FastFUCCI sensor contains constructs encoding mKO2-CDT1 to mark cells in G1, and mAG-geminin to mark cells in S/G2-M phases, both under the EF1α promoter^69^. We performed live imaging of A375-FUCCI cells under confinement to visualize how confinement affects the melanoma cell cycle. Upon applying confinement, all cells that were initially mitotic (mAG+, S/G2-M phase) rapidly lost mAG fluorescence, indicating they exited the cell cycle (**Fig. 5c-d**). Next, we knocked down HMGB2 using siRNA, which significantly impaired invasion of A375 human melanoma cells *in vitro* (*P* = 0.0133; **Fig. 5e-g**). To test the role of HMGB2 in phenotype switching *in vivo*, we generated *BRAF^V600E^*melanomas in zebrafish^71^ in which the zebrafish HMGB2 orthologs *hmgb2a* and *hmgb2b* were knocked out by CRISPR/Cas9. Loss of *hmgb2* and *hmgb2b* markedly increased melanoma growth, with HMGB2 CRISPR tumors growing almost 2-fold larger upon inactivation of hmgb2a/hmgb2b by 10 weeks (*P* = 0.0193; **Fig. 5h-j**). These data indicate that HMGB2 is required for the invasive state, and that its loss pushes melanoma cells towards a hyperproliferative state (**Fig. 6**).

**Figure 5:**
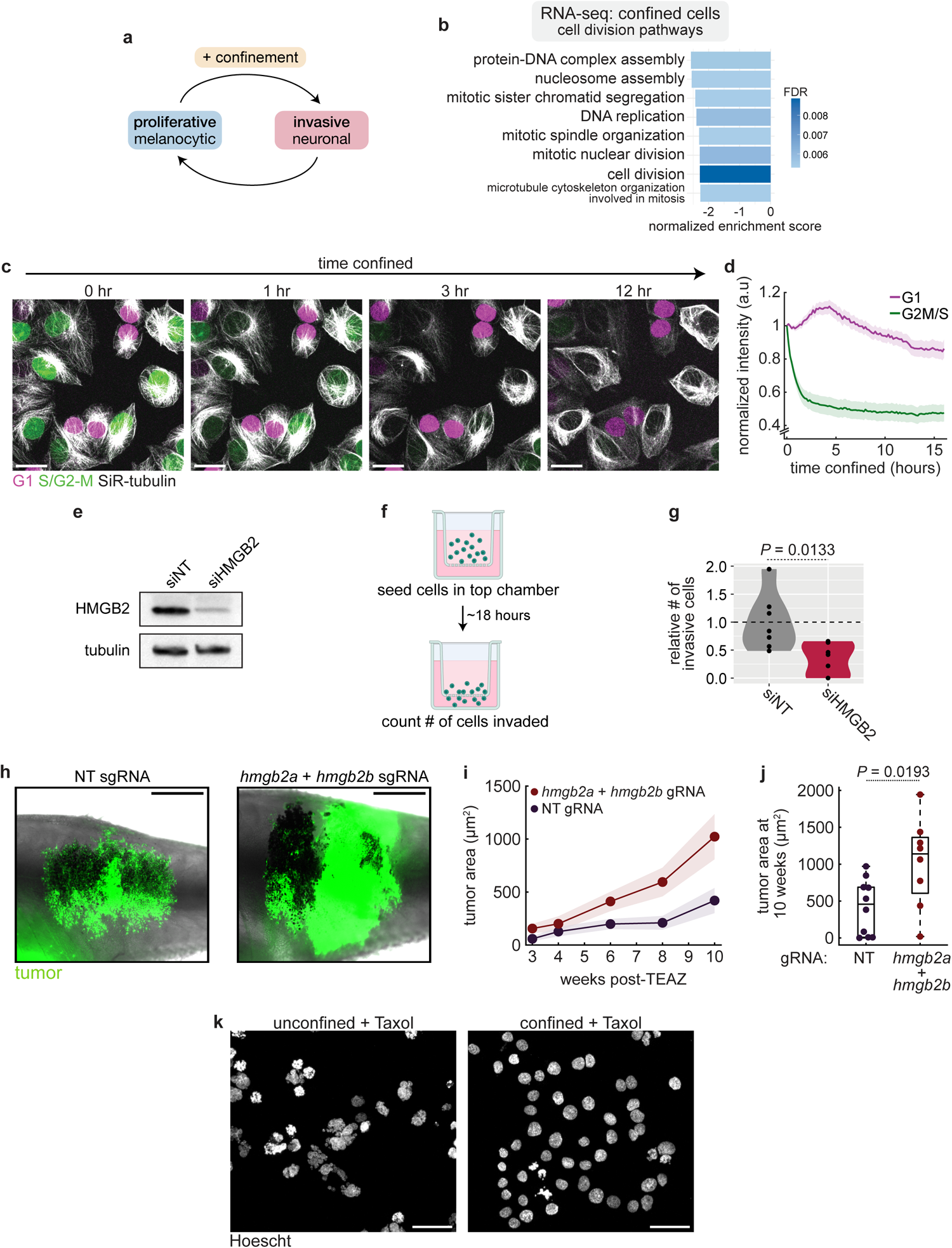
Confinement promotes drug tolerance by downregulating proliferation. **a.** Schematic detailing the influence of confinement on melanoma phenotype switching. **b.** Enrichment scores for GO:BP pathways related to cell division from RNA-seq of confined cells relative to unconfined. FDR is indicated. **c.** Stills from confocal imaging of A375 cells stably expressing the FASTFUCCI reporter and stained with SiR-tubulin. **d.** Quantification of FUCCI signal over time in confined cells. *n* = 37 cells from 4 movies. **e.** Western blot for HMGB2 (top) and tubulin (loading control, bottom) for A375 cells transfected with the indicated siRNAs. **f.** Schematic of in vitro invasion assay workflow. **g.** Quantification of *in vitro* invasion assay results for A375 transfected with the indicated siRNAs (*n* = 3 biological replicates). **h.** Representative images of adult zebrafish with melanomas generated using TEAZ, 12 weeks post electroporation. Tumor region is highlighted in red. Scale bars, 2 mm. **i.** Tumor surface area over time for melanomas induced in zebrafish using TEAZ. Error bars, SEM. **j.** Tumor surface area at 12 weeks post-TEAZ. **i-j.** sgNT: *n* = 20 fish. sgHMGB2: *n* = 16 fish. **k.** Hoescht (DNA) staining of cells treated with Taxol and confined (right) or unconfined control (left). Scale bars, 50 µm.

**Figure 6.**
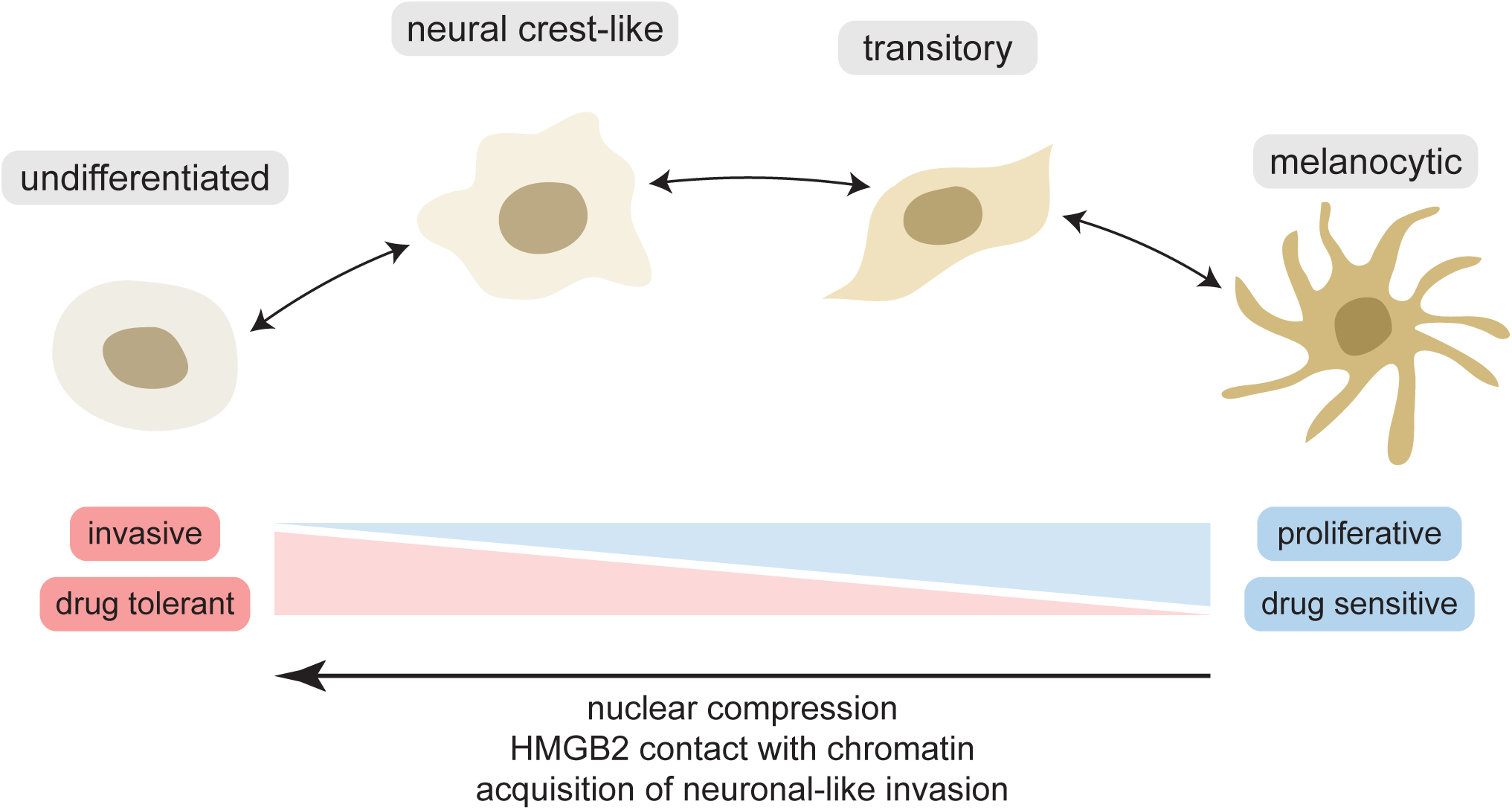
Confinement governs phenotypic plasticity in melanoma. Model for the role of mechanical confinement in melanoma phenotype switching: nuclear compression induces HMGB2 contact with chromatin, increasing chromatin accessibility and gene expression at neuronal loci.

Phenotype switching has been linked to drug resistance, with the undifferentiated/invasive cell state recently found to be drug tolerant^8,13,72^.This finding has important clinical relevance, as drug tolerant persister cells are a major contributor to relapse in patients^73^. Based on this, we examined how the confined, invasive state affects response to therapy. We hypothesized that the HMGB2-high neuronal state induced by confinement may contribute to drug tolerance in melanoma. To test this, we treated confined cells with Taxol to stabilize microtubules (**Fig. S5**). Whereas unconfined cells rapidly displayed a fragmented cell morphology and underwent apoptosis, the confined cells were almost completely resistant to Taxol-induced cell death (**Fig. 5k**). As a widely used chemotherapy, Taxol induces cancer cell death via cell cycle arrest^74^. This suggested that confinement may cause melanoma cells to exit the cell cycle, in accordance with a pro-invasive phenotype switch associated with downregulation of proliferation^14^ (**Fig. 6**).

## DISCUSSION

The emergence of single-cell sequencing has enabled the discovery of reproducible transcriptional states that correspond to distinct tumor phenotypes^6^. In melanoma, the extremes of these phenotype states encompass cells that are either highly proliferative or highly invasive. There is thought to be a tradeoff in these phenotypes, in which expression of the genes associated with proliferation (i.e. *MITF*) is mutually exclusive with those associated with invasion (i.e. *AXL*). This is reminiscent of the “go or grow” hypothesis that has been postulated to occur in both development and cancer, in which a cell is optimized for one phenotype over another^75,76^, similar to EMT^77^.

The reproducibility of these distinct states, in the absence of DNA lesions, has raised the possibility that they are epigenetically encoded. In our study, we find that the switch from a proliferative to invasive state is in part mediated by the chromatin-associated protein HMGB2. This family of proteins lacks specificity in their DNA-binding motifs, and instead form a protein complex with transcription factors or other cis-acting proteins. They are capable of bending DNA to facilitate the action of these other proteins^78^, making them ideal candidates for enforcing a particular chromatin configuration in response to external cues. Our data suggests that HMGB2 is responsive to the mechanical microenvironment. ATAC-seq analysis indicates that upregulation of HMGB2 is associated with a stable increase in accessibility at invasive loci, which may explain the persistence of the invasive state once a melanoma cell has encountered increased mechanical force from the microenvironment.

The role of the mechanical microenvironment on tumor cell phenotypes is a still emerging area^79^. Previous work has shown that increased pressure from a collagen matrix embedded with micro-beads could augment cancer cell invasion^80^, in part through changes in the actin cytoskeleton. Stromal cells such as cancer-associated fibroblasts can also change the mechanical microenvironment through secretion of MMPs that remodel that stiffness of the collagen matrix^81^. Our study adds to this field by showing that such mechanical force can also act through epigenetic remodeling of the cancer cell, which likely cooperates with signaling through mechanosensitive pathways such as YAP/TAZ/Hippo that promote metastatic colonization^82^. We did not find evidence of this pathway being activated in our system, suggesting there may be Hippo-independent mechanisms that respond to mechanical force in melanoma.

An interesting observation of our study is that mechanically confined melanoma cells enact an invasion program reminiscent of developing neurons. During development, neurons undergo passage through tightly confined spaces, and rearrange the tubulin network around the nucleus to protect it from high levels of force^40,41^, similar to what we observed in the invasive melanoma cells. An unexplored area is whether other aspects of neuronal signaling, aside from the acetylated tubulin cage, are important for melanoma invasion. Recent work from our lab has suggested that melanoma cells can use neuronal-like mechanisms during tumor initiation^83^, and future work should aim to explore this phenomenon in later steps of melanoma invasion.

## METHODS

### Zebrafish husbandry

Stable transgenic zebrafish lines were kept at 28.5°C in a dedicated aquatics facility with a 14 hour on/10 hour off light cycle. *casper* fish with the following genotype were used for all experiments: *mitfa-BRAF^V600E^; p53^-/-^; mitfa^-/-^*. Fish were anesthetized using Tricaine (MS-222; stock concentration 4g/L), diluted until the fish was immobilized. All animal procedures were approved by the Memorial Sloan Kettering Cancer Center IACUC (protocol #: 12-05-008).

### Zebrafish *in vivo* electroporation

Tumors were generated by Transgene Electroporation in Adult Zebrafish^71,84^ (TEAZ). To generate *hmgb2a*/*hmgb2b* knockout melanomas, adult (3-6 month old) fish were injected with the following plasmids: miniCoopR-GFP, mitfa:Cas9, Tol2, U6-sgptena, U6-sgptenb, and either 394-zU6-3XsgRNA[hmgb2a] and 394-zU6-3XsgRNA[hmgb2b], or 394-zU6-3XsgRNA[NT]. Adult fish were anesthetized in tricaine and injected with 1 µL of plasmid mixture below the dorsal fin, immediately electroporated, and moved to fresh water to recover. Tumor growth was imaged every 1-2 weeks using a Zeiss Axiozoom V16 fluorescence microscope.

### Cell culture

Human melanoma A375 cells were obtained from ATCC and maintained in a 37°C and 5% CO2 humidified incubator. Cells were routinely checked to be free from mycoplasma. Cells were cultured in DMEM (Gibco, 11965) supplemented with 10% FBS (Gemini Bio, 100-500).

### Cloning

To generate the HMGB2-GFP plasmid, the human HMGB2 coding sequence in a pENTR backbone (Horizon Discovery OHS5898-202621565) was combined with a C-terminus EGFP tag using InFusion cloning. The HMGB2-GFP insert was then transferred into a lentiviral expression vector containing the CMV promoter (pLX304) by Gateway cloning. To generate *hmgb2a* and *hmgb2b* CRISPR gRNA plasmids for use *in vivo*, 3 gRNAs for each gene were subcloned into Gateway entry vectors containing zebrafish-optimized U6 gRNA promoters. gRNAs were designed using CHOPCHOP^85^ and GuideScan^86^. The resulting 3X gRNA plasmid was assembled via Gateway LR cloning. gRNA cutting was validated *in vivo* using the Alt-R CRISPR-Cas9 system (IDT) by injecting sgRNAs and purified Cas9 protein into 1-cell stage zebrafish embryos. 24 hours later, genomic DNA was isolated from 5-10 embryos and mutation detection was performed using the Alt-R Genome Editing Detection Kit (IDT). gRNA sequences were:

*hmgb2a* sgRNA1: 5’-GAAAAGTTCACCGAGGTCCC-3’

*hmgb2a* sgRNA2: 5’-AAGGTGAAGGGCGACAACCC-3’

*hmgb2a* sgRNA3: 5’-GACAACCCGGGCATCTCTAT-3’

*hmgb2b* sgRNA1: 5’-CAAACCCAAGGGGAAGACGT-3’

*hmgb2b* sgRNA2: 5’-CTCAAACTTGACCTTGTCGG-3’

*hmgb2b* sgRNA3: 5’-AGAGAAGTTGACGGGCACGT-3’

NT sgRNA: 5’-AACCTACGGGCTACGATACG-3’

### Generation of stable cell lines

HMGB2^OE^, HMGB2-GFP and FastFUCCI stable A375 cell lines were generated by lentiviral transduction. The FastFUCCI reporter plasmid was obtained from Addgene (#86849). The HMGB2-GFP reporter plasmid was generated as described above. The HMGB2^OE^ plasmid was obtained from Horizon Discovery (OHS5897-202616132). 8 million HEK293T cells per condition were transfected with 1200 ng lentiviral vector, 600 ng PAX2 plasmid, and 300 ng MD2 plasmid, using Effectene transfection reagent (Qiagen). Virus was collected starting 24 hours after transfection. Viral supernatant was filtered (0.45 µm filter) before adding to A375 cells at a 1:1 ratio with media and 10 µg/mL polybrene. Cells were infected for 72 hours, allowed to recover for 24 hours, and then selected with blasticidin (5 µg/mL, 7 days) or puromycin (1 µg/mL, 3 days). To isolate brighter populations best for imaging, cells were then sorted using FACSAria III or FACSymphony S6 cell sorters (BD Biosciences). Expression of the untagged HMGB2 overexpression construct was validated using Western blotting with an antibody targeting HMGB2 (Millipore Sigma, HPA053314).

### In vitro confinement and imaging

A375 cells were subjected to overnight (∼16 hours) confinement at a height of 3 µm using a static cell confiner (4Dcell). Cells were plated 6 hours before imaging in fibronectin-coated glass-bottom 35mm dishes (Fluorodish) or glass-bottom 6-well plates (MatTek). Cells were allowed to attach before confinement was applied. Confined cells were incubated at 37°C and 5% CO_2_ overnight. For live imaging, dyes plus 10 µM verapamil were added to the plated cells 2-3 hours before imaging. Dyes used for live imaging were: SiR-tubulin (Spirochrome, 100 nM), SiR-DNA (Spirochrome, 250 nM). Pharmacological inhibitors were added immediately before applying confinement. Inhibitors used were: Taxol (Tocris 1097), tubacin (Selleck Chemicals S2239), nocodazole (Tocris 1228), and trichostatin A (Millipore Sigma T8552). Live imaging was performed on a LSM880 (Zeiss) confocal microscope at 37°C and 5% CO2, at 63X magnification and 5-10 minute temporal resolution, using Zen Black v2.3 SP1 software (Zeiss). For immunofluorescence, cells were fixed with 4% PFA for 15 mins at RT before proceeding with staining and imaging as described below.

### Immunofluorescence staining and imaging

Cells were plated on glass CC2-coated chamber slides (Thermo Fisher) or fibronectin-coated glass bottom dishes (Fluorodish) and allowed to attach for ∼24 hours. Cells were fixed with 4% PFA for 15 mins, permeabilized with 0.1% Triton in PBS, and blocked in 10% goat serum (Thermo Fisher) for 1 hour, all at RT. Primary antibodies used were: rabbit anti-HMGB2 (abcam, ab124670), rabbit anti-HMGB1 (abcam, ab18256), rabbit anti-HMGA1 (abcam, ab129153), mouse anti-α-tubulin (Millipore Sigma, CP06), chick anti-β-tubulin (Novus Biologicals, NB100-1612), mouse anti-acetylated tubulin (Millipore Sigma, 6793), rabbit anti-acetylated tubulin (Cell Signaling Technologies, CST 5335), rat anti-tyrosinated tubulin (Millipore Sigma, MAB1864-I), mouse anti-polyglutamylated tubulin (Millipore Sigma, T9822), mouse anti-GFP (abcam, ab1218), rabbit anti-H3Ac (Millipore Sigma, 06-599), mouse anti-Annexin V (Santa Cruz, sc-74438), rabbit anti-cleaved caspase-3 (CST 9661), rabbit anti-cleaved PARP (CST 5625). All primary antibodies were used at 1:200. Cells were incubated with primary antibodies overnight at 4°C, then washed in PBS and incubated with the appropriate fluorescently labelled secondary antibody (1:250). Alexa 488-conjugated phalloidin (CST 8878S), when used, was added at 1:50, and Hoescht added at 1:1000. Cells were mounted in Vectashield (Vector Laboratories) and allowed to cure overnight. Stained cells were imaged on a Zeiss LSM 880 confocal at 40X or 63X resolution, using Zen Black v2.3 SP1 software (Zeiss).

### Image analysis

Image analysis was done using CellProfiler^87^, TrackMate^88^, and MATLAB version R2021b and R2023b (Mathworks). For images of fixed cells, cells were segmented in CellProfiler using Hoescht staining to generate a nuclei mask, and phalloidin or other cytoskeletal staining to generate a whole-cell mask. The mean intensity per cell/nucleus was quantified, and expression of nuclear-localized proteins was normalized to Hoescht intensity per nucleus. For quantification of live imaging data, HMGB2-GFP intensity per cell over time was quantified using TrackMate. The resulting intensity data was analyzed in MATLAB, by fitting a line to each curve and automatically removing curves in which more than 4 data points differed from the line of best fit by more than 0.2 a.u. In all cases, plotting and statistics were done in MATLAB.

### Fluorescence recovery after photobleaching (FRAP)

A375 cells expressing HMGB2-GFP were confined for ∼18 hours before FRAP measurements. FRAP was done on a LSM 880 confocal at 37°C with 5% CO_2_, using a 63X oil immersion lens and Zen Black v2.3 SP1 software (Zeiss). A 5 µm circular diameter region of interest was defined within the nucleus of each cell, before photobleaching at 405 and 488 nm wavelengths for 10 pulses. One time point was acquired before photobleaching. Fluorescence recovery was imaged at 0.2 s intervals for a total of 20 s. All analysis was done in MATLAB. For analysis, fluorescence within the ROI was normalized to the fluorescence at the initial time point (pre-photobleaching). Samples in which the fluorescence within the ROI was not bleached to at least 25% of the pre-bleaching value were automatically removed from the analysis. Each recovery curve was fitted with a 2-component exponential using the function “fit” with the “exp2” parameter:

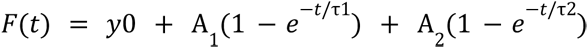

where *y0* represents the fluorescence immediately after photobleaching, *A_1_* represents the amplitude of the fast-diffusing population, *A_2_* represents the amplitude of the slow-diffusing population, *t* is time, and *τ1* and *τ2* correspond to the time constants for the fast- and slow-diffusing populations, respectively.

### Bulk RNA-sequencing and analysis

For bulk RNA-seq of A375 cells overexpressing HMGB2, 3 replicates of ∼1 million cells each were pelleted and resuspended in Trizol, before snap freezing. For bulk RNA-seq of confined A375 cells, 200,000 cells were plated in each well of a 6-well plate. 3 wells were confined for ∼18 hours using a 6-well static confiner (4Dcell) at 3 µm height while the remaining 3 wells were left unconfined. Cells were then collected in Trizol, pooling the 3 wells for each condition to generate samples of ∼600,000 cells each. This process was repeated for a total of 3 independent biological replicates per condition.

Library preparation and sequencing was done by Azenta Life Sciences. Raw sequencing reads were processed using FastQC (Babraham Bioinformatics) and Trimmomatic^89^ before alignment to the human genome hg38. All downstream analysis was performed in R (version 4.3.1). Differential gene expression analysis was performed using DESeq2^90^ with default parameters. Gene set enrichment analysis^91^ was done using the fgsea^92^ R package (version 1.26) with the GO^93,94^ biological processes pathway sets from MSigDB^95^.

### Bulk ATAC-sequencing and analysis

Samples containing ∼100,000 cells each were centrifuged at 700g for 5 min at 4°C before being resuspended in 500 µL growth media supplemented with 10% DMSO. Cells were frozen at −80°C overnight before library preparation and sequencing done by Azenta Life Sciences. Sequencing reads were trimmed and filtered for quality control using TrimGalore (version 0.6.7) with a quality setting of 15, cutadapt^96^ (version 4.0), and FastQC version 0.12.1. Reads were aligned to the human genome assembly hg38 using bowtie2^97^ (version 2.3.5.1) and were deduplicated using MarkDuplicates from Picard (Broad Institute; version 2.16). Peaks were identified using MACS2^98^ with a p-value setting of 0.001, using a publicly available melanocyte dataset (GSM3191792) as control. To generate a global peak atlas, blacklisted regions were removed, before merging all peaks within a 500 bp region and quantifying reads using featureCounts. Differentially enriched peaks were identified using DESeq2^90^. Peak-gene mapping was done by assigning all intergenic peaks to that gene, and in other cases by genomic distance to the transcription start site. Pathway analysis was done using clusterProfiler^99^. Tornado plots were generated with deepTools^100^ (version 3.5.1) functions computeMatrix and plotHeatmap with genes annotated from the indicated pathway sets. Motif enrichment analysis was performed using HOMER^60^ (version 4.11.1) functions findMotifsGenome and annotatePeaks.

### Re-analysis of human melanoma scRNA-seq data

Human melanoma scRNA-seq data from ref.^12^ was downloaded from GEO (GSE115978). All analyses were performed in R using Seurat^101^ version 4.4.0 and 5.0.1. The counts matrix was normalized using SCTransform^102^. Clustering was done using the Seurat functions FindNeighbors and FindClusters with a resolution of 0.8. Cell types and treatment status were annotated using metadata from the original publication^12^. Cell types were classified using gene lists from ref.^13^ and the Seurat function AddModuleScore with default parameters. Module scores were scaled between 0 and 1. Cells were classified by differentiation state based on the highest expression score for the given gene modules. Differentially expressed genes were calculated using the Seurat function FindMarkers with default parameters. GSEA was performed using fgsea as described above.

### Statistical analysis

All statistical analysis and plotting were performed in either R (for RNA-seq and ATAC-seq data, version 4.3.1) or MATLAB (for imaging data, version R2021b). For scRNA-seq data, *P* values were calculated using the Wilcoxon rank sum test with Bonferroni’s correction for multiple groups (R functions pairwise.wilcox.test). Pearson correlation coefficients and corresponding *P* values were calculated using the R function cor.test. For differential expression analysis of bulk RNA-seq and bulk ATAC-seq data, *P* values were calculated in DESeq2 using the Wald test. For image analysis, *P* values were calculated using MATLAB functions anova1 and multcompare using the Tukey post-hoc test.

## Data availability

Raw and processed RNA-seq and ATAC-seq data generated in this study have been deposited to the Gene Expression Omnibus (GEO) under accession number GSE253803. Human melanoma scRNA-seq data was obtained from GEO accession number GSE115978. All other relevant data supporting the key findings of this study are available within the article and its Supplementary Information files or from the corresponding authors upon request.

## Code availability

All code used for analysis and plotting is available at github.org/mvhunter1/Hunter_2024.

## ACKNOWLEDGEMENTS

We thank the members of the White lab for useful discussions. We thank the MSKCC Molecular Cytology core facility for assistance with imaging, the MSKCC Aquatics core facility for zebrafish care and maintenance, and the MSKCC Flow Cytometry core facility for assistance with FACS. MVH was funded by a K99/R00 Pathway to Independence Award from the National Cancer Institute (1K99CA266931), the Scholarship for the Next Generation of Scientists from the Cancer Research Society, and a postdoctoral fellowship from the Canadian Institutes of Health Research. YM was supported by a Medical Scientist Training Program grant from the NIH under award number T32GM007739 to the Weill Cornell/Rockefeller/Sloan Kettering Tri-Institutional MD-PhD Program and Kirschstein-National Research Service Award (NRSA) predoctoral fellowship under award number F30CA265124. IY and RMW were funded by NIH Research Program Grant 1U01CA260432. RMW was funded by the Melanoma Research Alliance, The Debra and Leon Black Family Foundation, NIH Research Program Grants R01CA229215 and R01CA238317, the NIH Director’s New Innovator Award DP2CA186572, the Pershing Square Sohn Foundation, The Mark Foundation, The Alan and Sandra Gerry Metastasis Research Initiative at MSKCC, The Harry J. Lloyd Foundation, Consano, the Starr Cancer Consortium, and the American Cancer Society RSG-19-024-01-DDC.

## AUTHOR CONTRIBUTIONS

MVH and RMW conceived the study. MVH performed all experiments and analyses unless otherwise noted. EM and YM performed TEAZ experiments. RM and IY assisted with transcriptomics experiments. RPK assisted with ATAC-seq analysis. MVH and RMW wrote the manuscript and all authors provided feedback before submission.

## MATERIALS AND CORRESPONDENCE

All correspondence and requests for materials should be directed to Miranda V. Hunter or Richard M. White.

