## Supplementary Figures for "Mechanical confinement governs phenotypic plasticity in melanoma"

### Figure S1

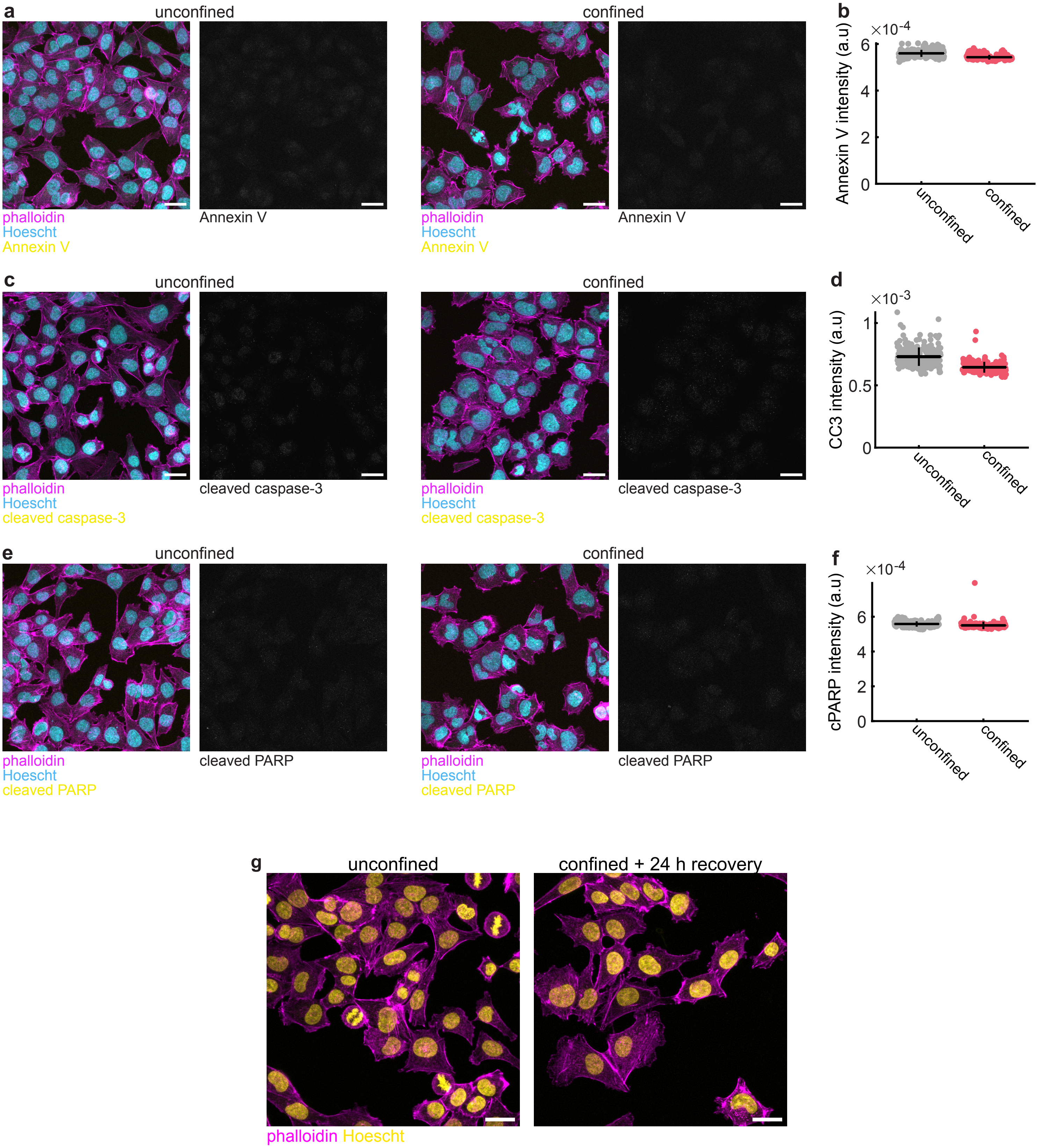

a
unconfined
confined
phalloidin
Hoescht
Annexin V
Annexin V
b
×10-4
Annexin V intensity (a.u)
6
4
2
0
unconfined
confined
c
unconfined
confined
phalloidin
Hoescht
cleaved caspase-3
cleaved caspase-3
d
×10-3
CC3 intensity (a.u)
1
0.5
0
e
unconfined
confined
phalloidin
Hoescht
cleaved PARP
cleaved PARP
f
×10-4
cPARP intensity (a.u)
g
unconfined
confined + 24 h recovery
phalloidin Hoescht

**Figure S1: Confinement does not cause apoptosis. a,c,e.** IF of confined A375 cells labelled with antibodies against the apoptosis markers Annexin V (a), cleaved caspase-3 (c), and cleaved PARP (e). **b,d,f.** Intensity per cell for the indicated markers. **b.** Unconfined: $n$ = 250 cells from 6 images. Confined: $n$ = 225 cells from 6 images. **d.** Unconfined: $n$ = 261 cells from 6 images. Confined: $n$ = 185 cells from 6 images. **f.** Unconfined: $n$ = 245 cells from 6 images. Confined: $n$ = 174 cells from 6 images. **g.** IF of A375 cells confined for ~18 hours and then left to recover for 24 hours (right) or unconfined cells (right). **a,c,e,g.** Scale bars, 25 µm.

### Figure S2

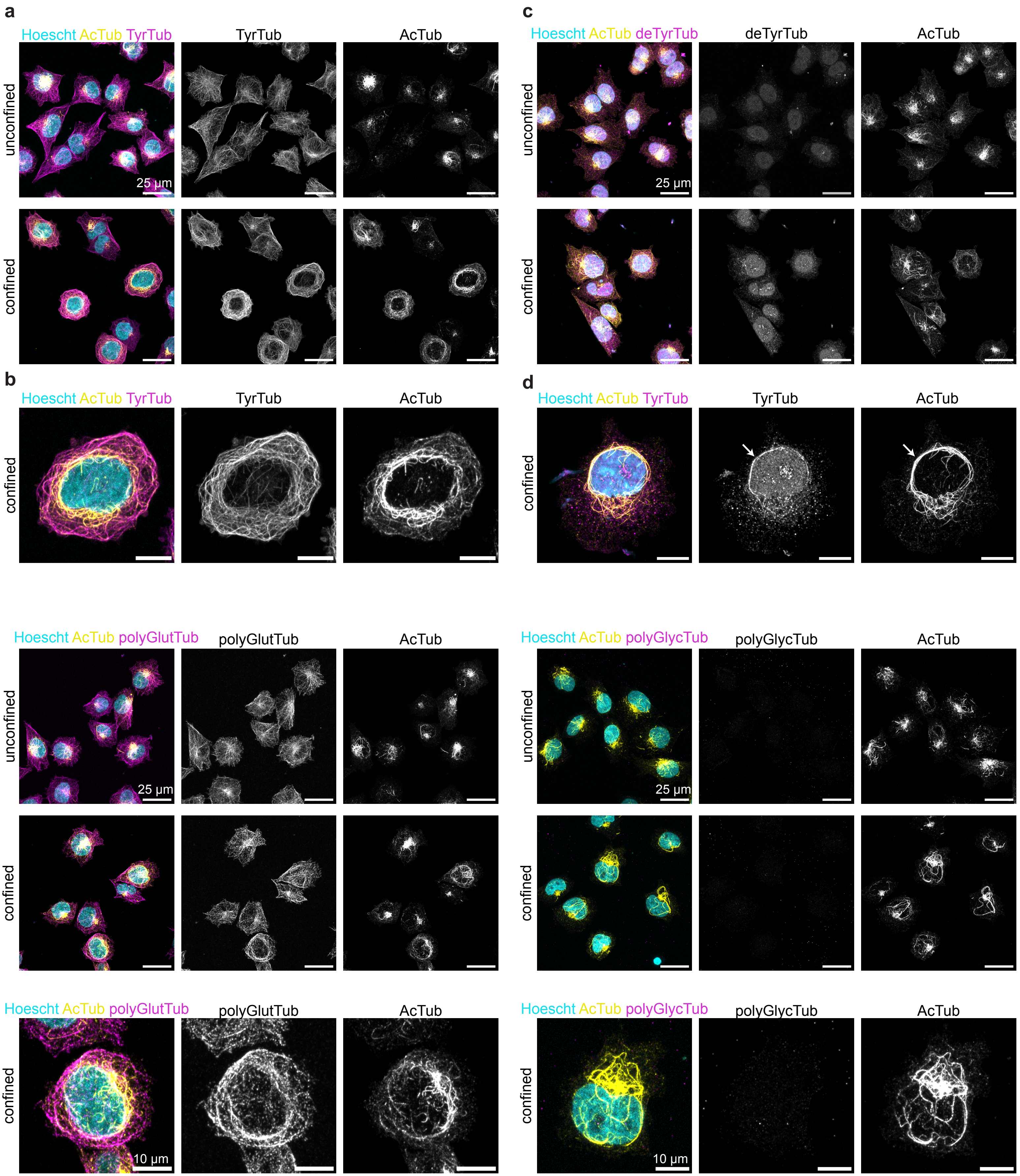

a
Hoescht AcTub TyrTub
TyrTub
AcTub
unconfined
confined
25 µm
c
Hoescht AcTub deTyrTub
deTyrTub
AcTub
unconfined
confined
25 µm
b
Hoescht AcTub TyrTub
TyrTub
AcTub
confined
d
Hoescht AcTub TyrTub
TyrTub
AcTub
confined
Hoescht AcTub polyGlutTub
polyGlutTub
AcTub
unconfined
confined
25 µm
Hoescht AcTub polyGlycTub
polyGlycTub
AcTub
unconfined
confined
25 µm
Hoescht AcTub polyGlutTub
polyGlutTub
AcTub
confined
10 µm
Hoescht AcTub polyGlycTub
polyGlycTub
AcTub
confined
10 µm

**Figure S2: Localization of tubulin post-translational modifications in confined cells. a-h.** IF of A375 cells labeled with antibodies targeting acetylated tubulin, in addition to tyrosinated tubulin (a-b), detyrosinated tubulin (c-d), polyglutamylated tubulin (e-f), and polyglycylated tubulin (g-h). **a,c,e,g.** Scale bars, 25 µm. **b,d,f,h.** Scale bars, 10 µm.

#### Figure S3

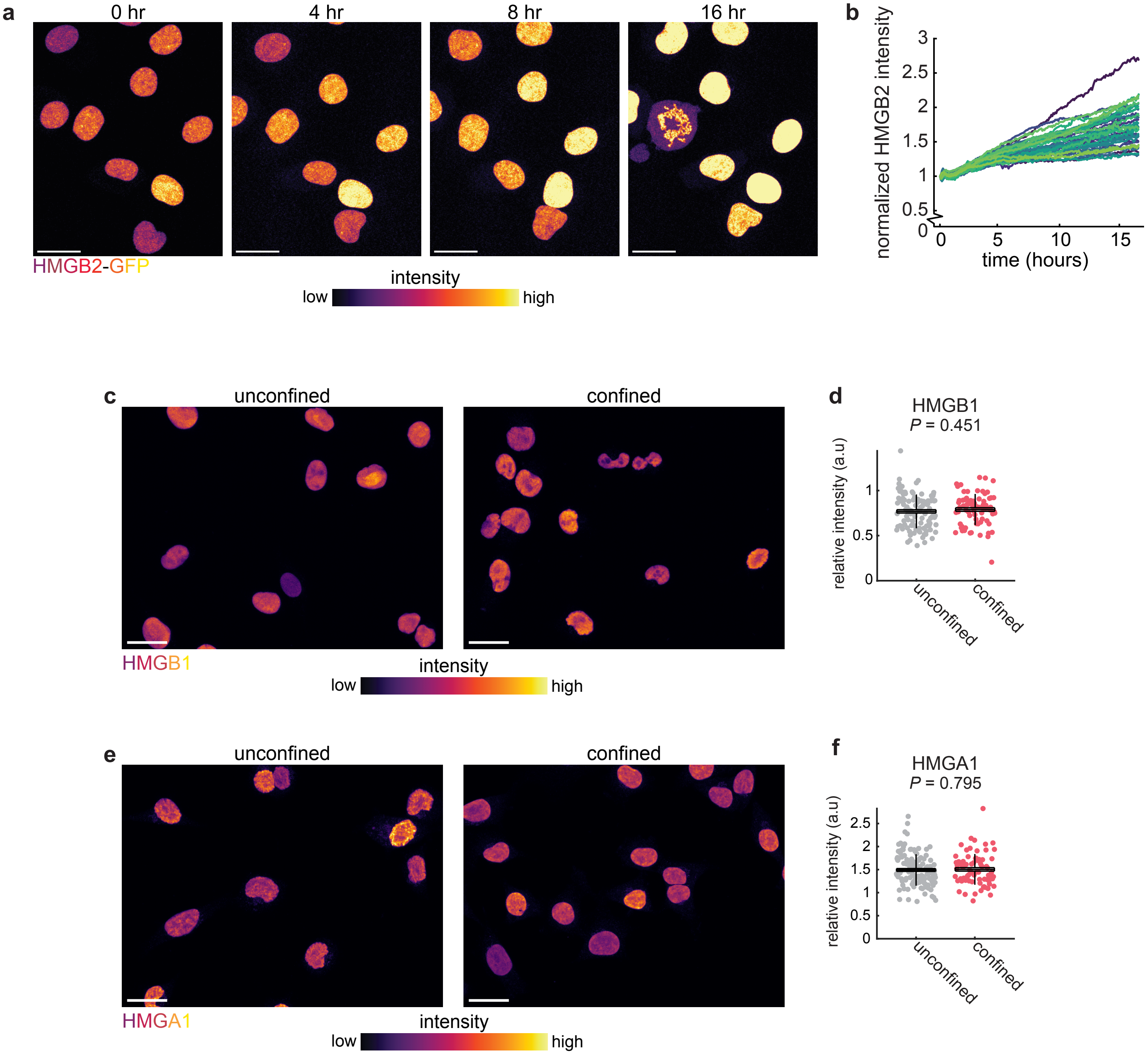
a
0 hr
4 hr
8 hr
16 hr
HMGB2-GFP
intensity
low
high
b
normalized HMGB2 intensity
0
0.5
1
1.5
2
2.5
3
0
5
10
15
time (hours)
c
unconfined
confined
HMGB1
intensity
low
high
d
HMGB1
P = 0.451
relative intensity (a.u)
0
0.5
1
unconfined
confined
e
unconfined
confined
HMGA1
intensity
low
high
f
HMGA1
P = 0.795
relative intensity (a.u)
0
0.5
1
1.5
2
2.5
unconfined
confined

**Figure S3: Confinement specifically upregulates HMGB2. a.** Stills from confocal imaging of A375 cells expressing HMGB2-GFP. Cells are pseudocolored by HMGB2 intensity. **b.** HMGB2-GFP intensity per cell over time, normalized to intensity at the first time point acquired. $n$ = 37 cells from 6 movies. **c,e.** Immunofluorescence images of A375 cells stained with antibodies targeting HMGB1 (c) and HMGA1 (e). Scale bars, 25 µm. **d,f.** Quantification of intensity in confined/unconfined cells for the indicated markers. Each point represents 1 cell. Horizontal lines, mean; box, SEM; vertical lines, SD. **d.** Unconfined: $n$ = 105 cells from 9 images. Confined: $n$ = 76 cells from 9 images. **f.** Unconfined: $n$ = 125 cells from 9 images. Confined: $n$ = 80 cells from 9 images.

### Figure S4

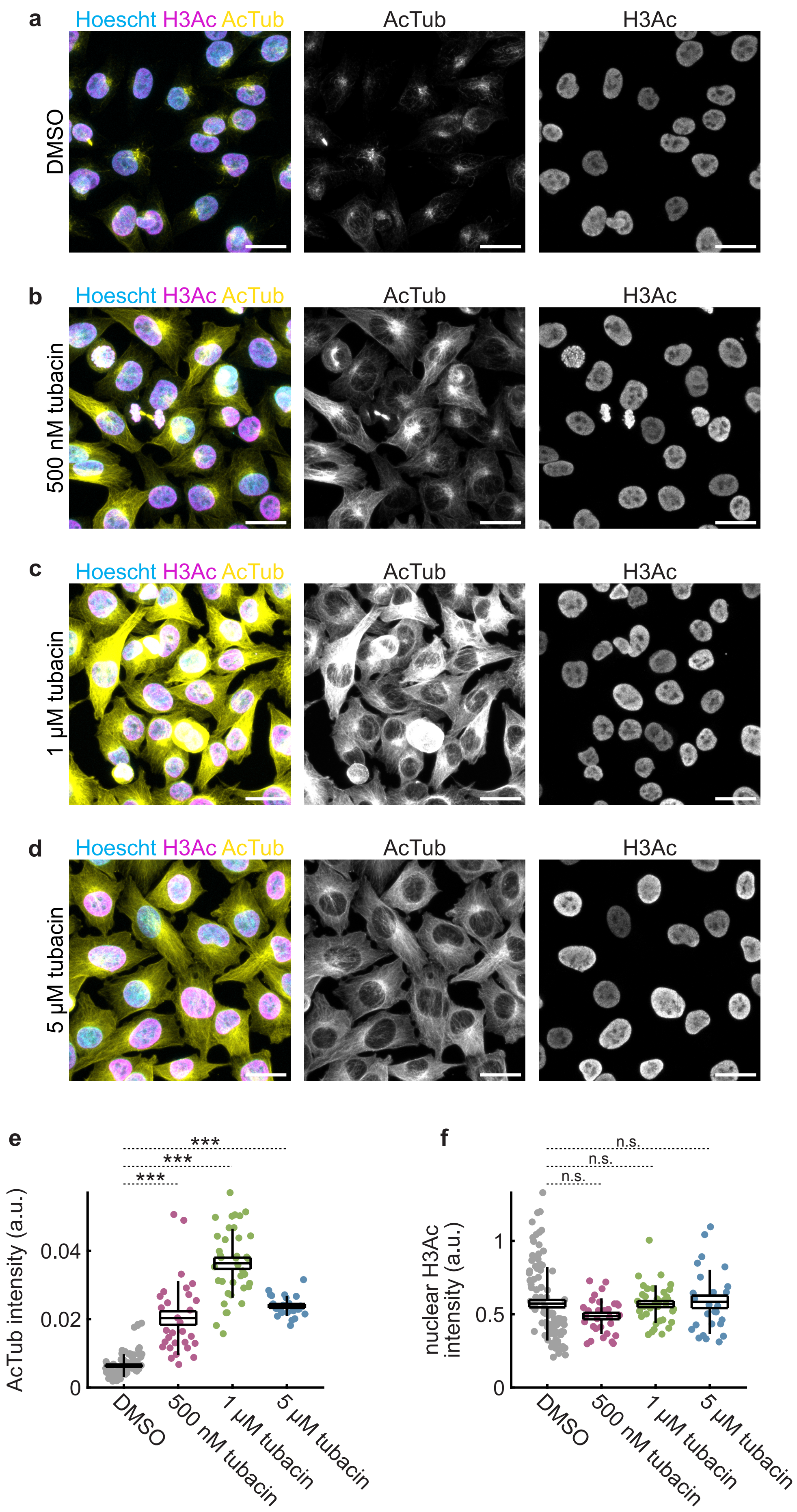
a
Hoescht H3Ac AcTub
AcTub
H3Ac
DMSO
b
500 nM tubacin
c
1 µM tubacin
d
5 µM tubacin
e
***
***
***
AcTub intensity (a.u.)
0.04
0.02
0
DMSO
500 nM tubacin
1 µM tubacin
5 µM tubacin
f
n.s.
n.s.
n.s.
nuclear H3Ac intensity (a.u.)
1
0.5
0

**Figure S4: Tubacin induces tubulin hyperacetylation without affecting histone acetylation. a-d.** A375 cells treated with the indicated tubacin concentrations (b-d) or DMSO as a vehicle control (a) and stained with antibodies labelling acetylated histone H3 and acetylated tubulin. Scale bars, 25 µm. **e-f.** Quantification of whole-cell acetylated tubulin intensity (e) and nuclear histone H3 acetylation (g). ***, $P < 0.001$. n.s., not significant.

### Figure S5

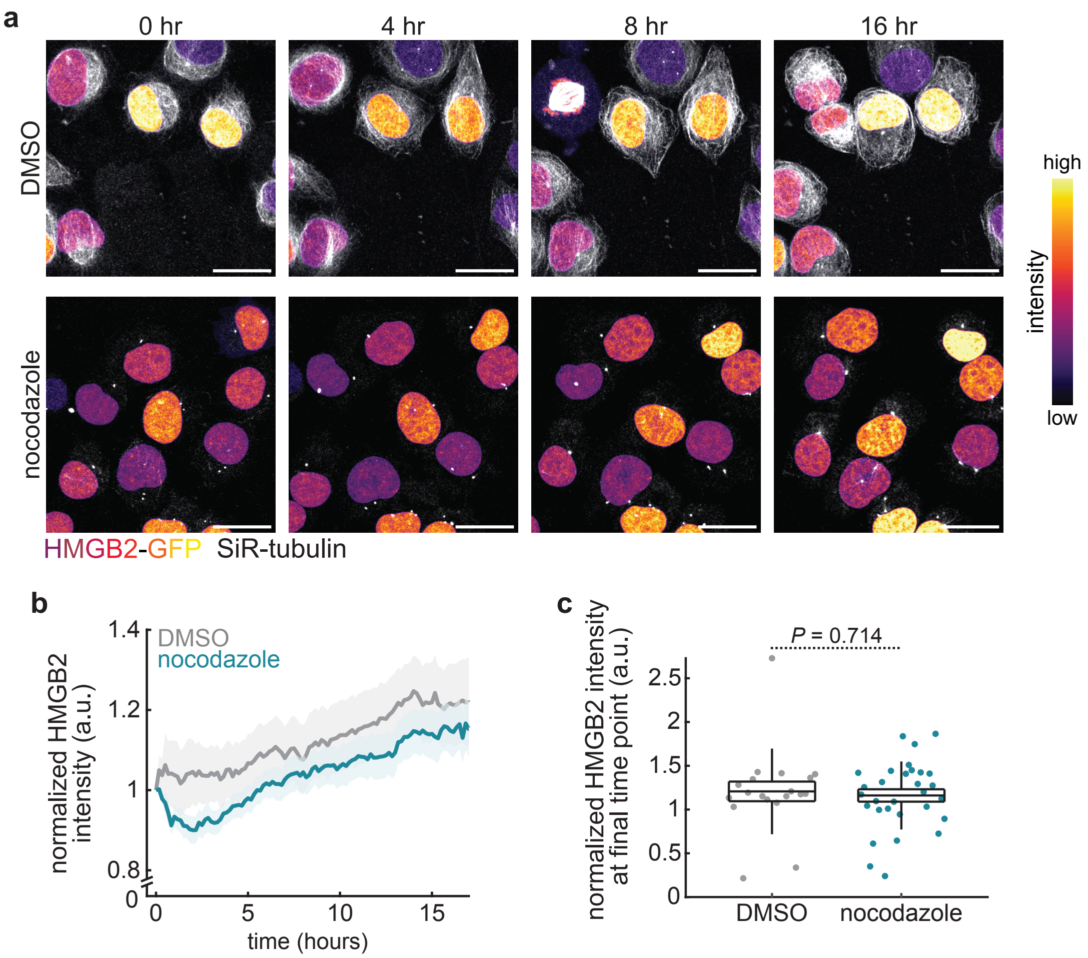

a
0 hr
4 hr
8 hr
16 hr
DMSO
nocodazole
high
intensity
low
HMGB2-GFP SiR-tubulin
b
normalized HMGB2 intensity (a.u.)
1.4
1.2
1
0.8
0
DMSO
nocodazole
0
5
10
15
time (hours)
c
normalized HMGB2 intensity at final time point (a.u.)
P = 0.714
2.5
2
1.5
1
0.5
0
DMSO
nocodazole

**Figure S5: Tubulin stabilization upregulates nuclear HMGB2. a.** HMGB2-GFP and SiR-tubulin intensity over time for confined cells treated with DMSO or 1 µM nocodazole. Scale bars, 25 µm. **b.** HMGB2-GFP intensity over time. Error bars, SEM. **c.** HMGB2-GFP intensity per cell at the final time point imaged (~ 16 hours). Horizontal lines, mean; box, SEM; vertical lines, SD. **b-c.** DMSO: *n* = 19 cells from 3 movies. Nocodazole: *n* = 30 cells from 3 movies.

### Figure S6

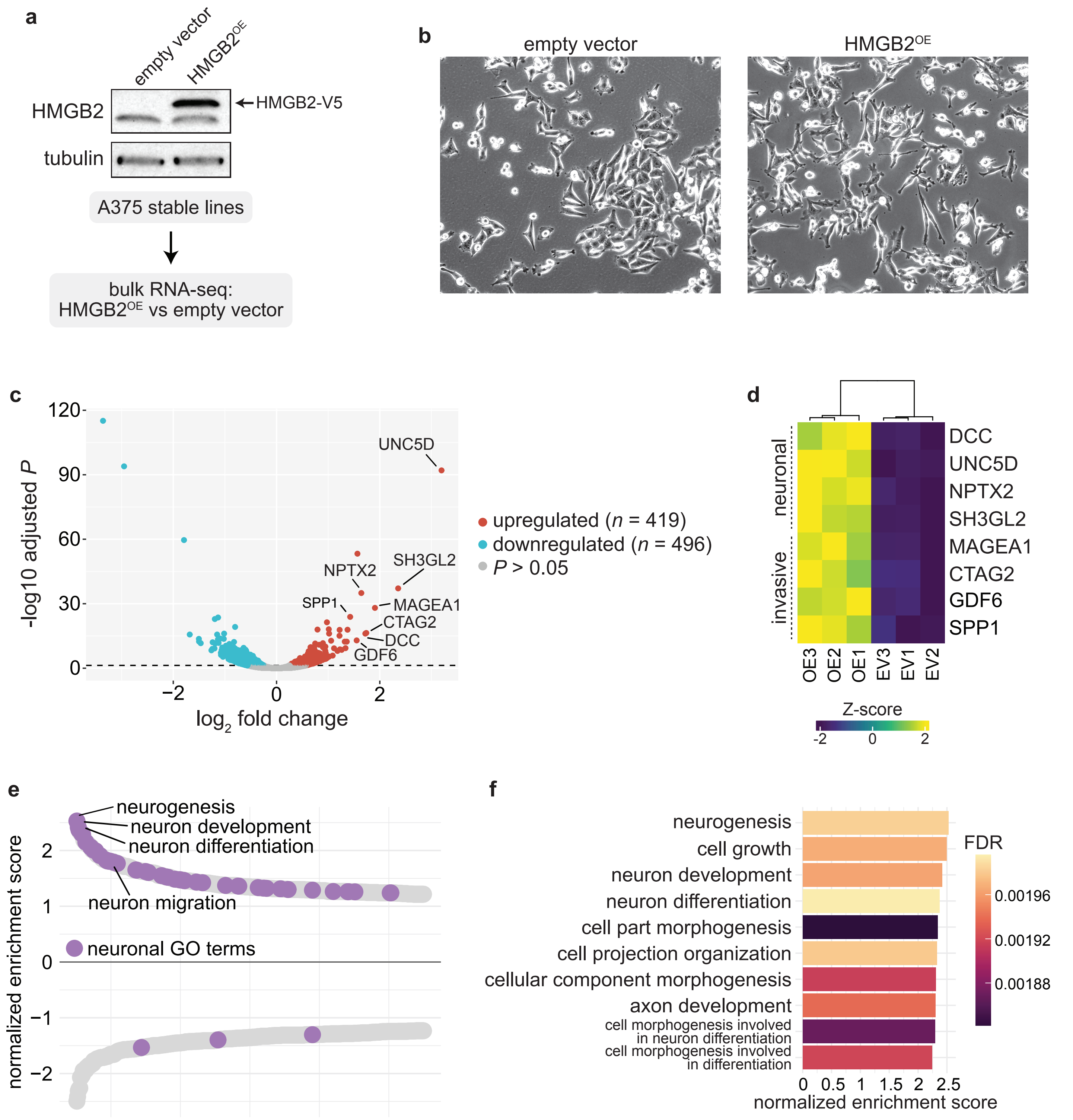
a
empty vector
HMGB2OE
HMGB2
HMGB2-V5
tubulin
A375 stable lines
bulk RNA-seq:
HMGB2OE vs empty vector
b
empty vector
HMGB2OE
c
UNC5D
NPTX2
SH3GL2
SPP1
MAGEA1
CTAG2
DCC
GDF6
120
90
60
30
0
-log10 adjusted P
−2
0
2
log2 fold change
upregulated (n = 419)
downregulated (n = 496)
P > 0.05
d
neuronal
invasive
DCC
UNC5D
NPTX2
SH3GL2
MAGEA1
CTAG2
GDF6
SPP1
OE3
OE2
OE1
EV3
EV1
EV2
Z-score
-2
0
2
e
neurogenesis
neuron development
neuron differentiation
neuron migration
neuronal GO terms
normalized enrichment score
2
1
0
−1
−2
f
neurogenesis
cell growth
neuron development
neuron differentiation
cell part morphogenesis
cell projection organization
cellular component morphogenesis
axon development
cell morphogenesis involved in neuron differentiation
cell morphogenesis involved in differentiation
FDR
0.00196
0.00192
0.00188
0
0.5
1
1.5
2
2.5
normalized enrichment score

**Figure S6: HMGB2 overexpression induces a neuronal gene program. a.** Western blot for HMGB2 (top) and tubulin (loading control, bottom) in stable A375 lines infected with lentivirus encoding HMGB2 or an empty vector control. **b.** Representative images of cells from cell lines indicated in a. **c.** Volcano plot of differentially expressed genes upon HMGB2$^{OE}$. **d.** Heatmap of Z-scored expression of selected neuronal and invasive genes across replicates. **e.** Double waterfall plot of top GO biological processes pathways by normalized enrichment score (NES). Neuronal pathways are labeled in purple. **f.** Top 10 GO biological processes pathways by NES upregulated upon HMGB2$^{OE}$.
